# An improved model for prediction of de novo designed proteins with diverse geometries

**DOI:** 10.1101/2025.06.02.657515

**Authors:** Benjamin Orr, Stephanie E. Crilly, Deniz Akpinaroglu, Eleanor Zhu, Michael J. Keiser, Tanja Kortemme

## Abstract

Nature uses structural variations on protein folds to fine-tune the geometries of proteins for diverse functions, yet deep learning-based de novo protein design methods generate highly regular, idealized protein fold geometries that fail to capture natural diversity. Here, using physics-based design methods, we generated and experimentally validated a dataset of 5,996 stable, de novo designed proteins with diverse non-ideal geometries. We show that deep learning-based structure prediction methods applied to this set have a systematic bias towards idealized geometries. To address this problem, we present a fine-tuned version of Alphafold2 that is capable of recapitulating geometric diversity and generalizes to a new dataset of thousands of geometrically diverse de novo proteins from 5 fold families unseen in fine-tuning. Our results suggest that current deep learning-based structure prediction methods do not capture some of the physics that underlie the specific conformational preferences of proteins designed de novo and observed in nature. Ultimately, approaches such as ours and further informative datasets should lead to improved models that reflect more of the physical principles of atomic packing and hydrogen bonding interactions and enable improved generalization to more challenging design problems.

## INTRODUCTION

State-of-the-art deep learning models such as AlphaFold2^1^ (AF2), AlphaFold3^2^ (AF3), RoseTTAFold (RF)^3^, and ESMFold^4^ are able to predict the structure of proteins from sequence with atomic accuracy in many cases. These advances have led to considerable success with the inverse problem of structure prediction: protein design, the generation of proteins with new structures and functions^5^. A general workflow involves, first, sampling a protein backbone using a generative method, such as RFdiffusion^6^ or AF2-based hallucination^7,8^, and second, predicting sequences optimal for these backbones using graph neural networks, such as ProteinMPNN^9^, or structure-conditioned language models, such as Frame2seq^10^. In an additional step, candidate designs are typically validated in silico for whether the designed sequences fold into their intended structures using models such as AF2^1^ or ESMFold^4^. Designs passing these in silico filters are then experimentally tested, and reported success rates for experimentally validated de novo protein binders to hydrophobic target sites can be as high as 10% or greater^8^.

The increasing number of experimentally validated de novo designed proteins now provides opportunities to compare the sequence and structure space sampled by de novo proteins generated with deep learning to that sampled by naturally occurring, evolved proteins^11,12^. Several biases have become apparent. An early observation was the vast overrepresentation of all-helical structures in de novo designed proteins^13,14^. Moreover, current examples of de novo-designed *functional* proteins are almost exclusively all-helical^14^. While methods can be conditioned to instead produce alpha-beta or all-beta proteins, successful examples have been relatively scarce. A second key observation is that de novo proteins generated by deep learning methods tend to adopt primarily idealized structures that lack features present in most evolved proteins, such as irregular secondary structure orientations and long loops with unique conformations^12^.

Such deviations from highly regular geometries may however be necessary to accommodate the precise orientations of chemical groups in protein active sites required for function^15,16^. While idealized de novo proteins are often extremely stable^17^, engineering function into them is still challenging. Despite increasing success designing protein binders targeting hydrophobic surfaces^6,8,18^, success rates for designing the loop-rich interfaces of antibody-antigen interactions^19^, binding sites for small molecules^20,21^, and constellations of catalytic groups in enzyme active sites^6^ are often low. This is likely because it is difficult to satisfy the geometric requirements of these functional sites with idealized arrangements of secondary structures. Indeed, naturally occurring proteins achieve functional site geometries not by using a different idealized fold topology for each function, but typically by diversifying the geometries of structural elements within a given fold^22^. To address this issue, several methods have been developed to mimic the ability of natural proteins to sample variations in sizes, orientations, and positions of secondary structure elements to generate geometries optimal for function. For example, structure prediction and design methods using the Rosetta software suite have been adapted to systematically sample variations on common natural alpha-beta folds, such as the Rossmann fold family, which is overrepresented in enzymes^22^, the NTF2 family, which is prevalent in ligand binding proteins^22,23^, and other folds^24^.

Here we ask whether these tunable geometric variations of common alpha-beta protein folds – which were previously engineered successfully using physics-based methods^22^ – would be predicted and designed with newer deep learning methods. We find that for a large set of stable de novo designed proteins with diverse geometries, predictions from AF2 substantially deviate from the physics-based design models. Moreover, we show that the AF2 predictions are systematically biased towards structures that were previously designed to have idealized geometries^17,25^. To test whether this problem can be alleviated, we fine-tune AF2 on a dataset of ∼6,000 stable proteins designed to adopt diverse geometries of the Rossmann fold. We show that the fine-tuned model predicts structures with improved sequence-structure compatibility for geometrically diverse de novo proteins and generalizes to other alpha-beta folds not used in fine-tuning. Improved prediction and design of non-idealized geometries by such a fine-tuned model should improve the design of more advanced functions requiring precise functional group positioning in active and polar binding sites.

## RESULTS

Naturally occurring proteins achieve the precise active site geometries necessary for function by diversifying the geometries of structural elements within a given fold. We set out to test to what extent deep learning methods used in state-of-the-art design workflows are capable of mimicking this behavior of natural fold families. We first tested the ability of a state-of-the-art de novo backbone generation method, RFdiffusion^6^, and a previously described Rosetta-based method, LUCS^22^, to generate geometric diversity in a common protein fold, the 2x2 Rossmann topology. To do so, we used both methods to diversify the positions, orientations, and sizes of two helices and their connecting loops on one side of the central beta sheet as a test case (**Fig. 1A**), because these loop-helix-loop (LHL) elements had been experimentally validated in a previous de novo design study to be capable of adopting diverse geometries^22^. We generated thousands of structurally diverse backbones with both LUCS^22^ and RFdiffusion^6^, sampling structural diversity in the same two LHL elements. We then compared the average pairwise backbone root mean square deviations (RMSDs) of the generated helix geometries to those present in 44 natural members of the Rossmann fold family (Methods, **Supplementary Fig. 1**, **Supplementary Data 1**). We find that LUCS generates helix geometries (6.8 Å average pairwise helix RMSD) that approach the structural diversity of natural Rossmann folds (6.9 Å average pairwise RMSD), and that both surpass the much more limited structural diversity generated by RFdiffusion (4.7 Å average pairwise RMSD) (**Fig. 1B**). Visualizing structural examples from each of the three sets illustrates that RFdiffusion generates more regular geometries (**Fig. 1C**) with primarily straight helices parallel to the underlying beta strands, in contrast to the more varied geometries in both the natural and LUCS-generated structures (**Fig. 1D**). These results suggest that new deep learning methods focused specifically on diversifying structural geometries may be required to capture and design geometric diversity similar to that observed in nature^12^.

**Figure 1.**
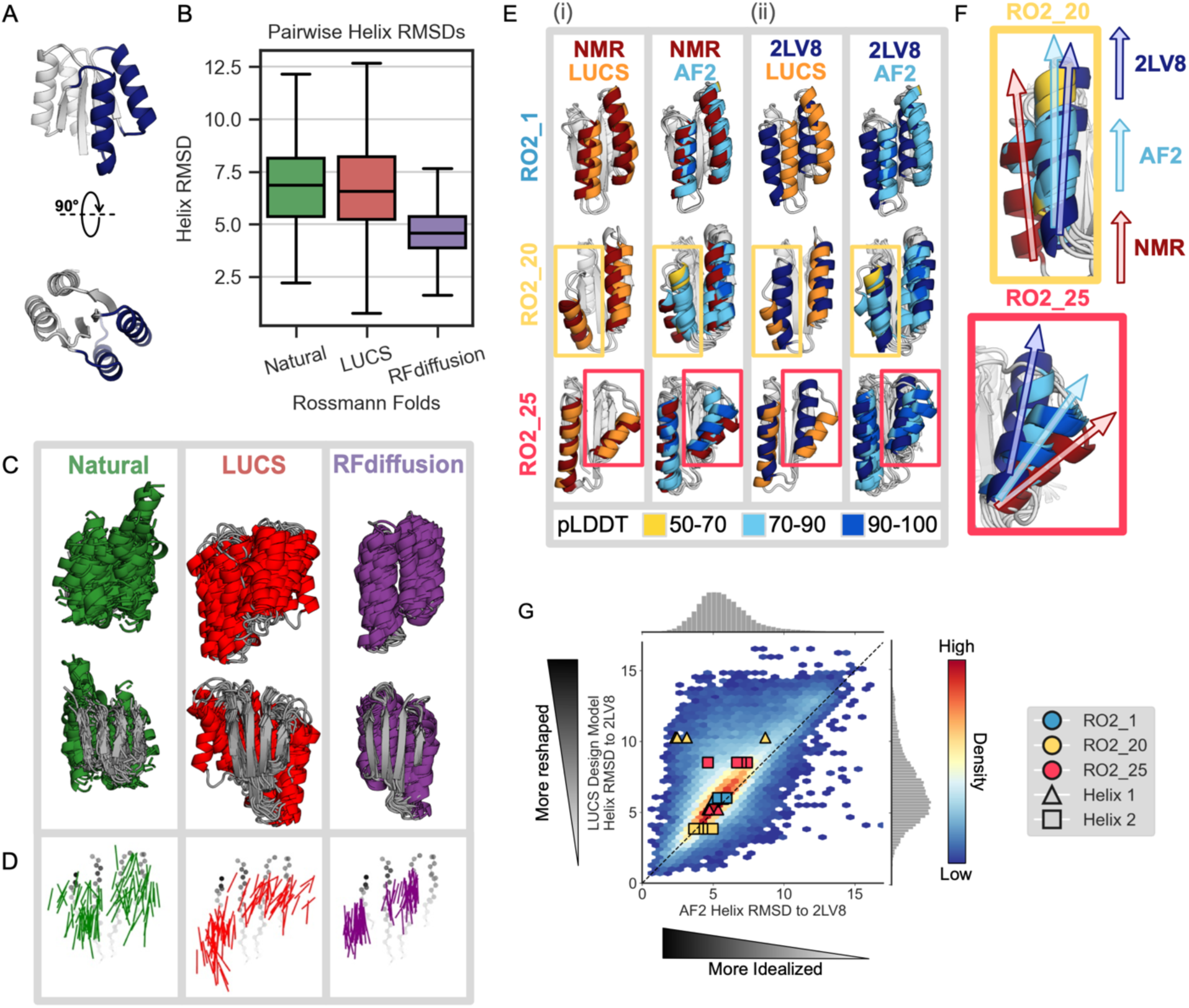
Deep learning-based protein structure prediction and design models predict and generate more idealized geometries than those occurring in nature or in de novo proteins designed using physics-based methods. **(A)** Idealized Rossmann fold (PDB ID 2LV8) with loop-helix-loop (LHL) units diversified in this study shown in dark blue. **(B)** Comparison of geometric variation within groups of Rossmann fold backbones. Natural (green): 44 representative natural Rossmann fold proteins. LUCS (red): 5,996 backbones generated with loop-helix-loop unit combinatorial sampling^22^. These designed backbones were experimentally determined to fold into stable proteins (Fig. 2). RFdiffusion (purple): 5,996 backbones generated with RFdiffusion^6^ (Methods). Boxes show the lower and upper quartiles, whiskers extend to points within 1.5 interquartile ranges of the lower and upper quartiles, and lines indicate the median values (outliers not shown). **(C)** Structural overlay of 44 examples from each Rossmann fold group. The 44 LUCS and RFdiffusion examples were selected by sorting backbones by their helix RMSDs to an idealized Rossmann fold (2LV8^17^), then sampling at even intervals to span the helix RMSD range. **(D)** Vector representations of helices to depict geometric diversity, with the underlying beta-sheet shown in grey. Vectors indicate the helix centroids (vector tails) and average carbonyl C-O direction. **(E)** Structural comparisons for three experimentally validated LUCS designs (RO2_1, RO2_20 and RO2_25 from ref.^22^) with reshaped helices shown in color: LUCS design models (orange) and 5 AF2 predictions (colored by pLDDT) are compared to (i) their experimentally-determined lowest energy NMR structures (dark red, showing close agreement with LUCS), and (ii) an idealized 2x2 Rossmann fold de novo designed protein (dark blue; PDB ID: 2LV8^17^). **(F)** Closeup of comparison for non-idealized helices in RO2_20 and RO2_25, showing that AF2 predicts helix geometries closer to the idealized structure than the experimentally determined structure. **(G)** Comparison of AF2 structure predictions to the LUCS design models for 10,000 LUCS backbones with sequences designed with ProteinMPNN. AF2 predicts 75.8% of reshaped helices to be closer to the idealized helices in 2LV8 than to the LUCS design model (density above the diagonal). Overlaid points show the helix RMSDs to 2LV8 for the design models and AF2 predictions for the three designs with experimentally determined structures in (E) and (F).

We next asked whether the limited diversity generated by RFdiffusion is at least in part due to limitations of current deep learning methods for structure prediction such as AF2 or RF (on which backbone generation methods such as AF2-hallucination or RFdiffusion are based) to generalize to protein backbones with non-ideal structural geometries. We first evaluated deep learning-based structure predictions for three de novo designed 2x2 Rossmann fold proteins with experimentally determined structures that have geometries different from the idealized 2x2 Rossmann fold protein (PDB ID 2LV8) originally designed by Koga et al.^17^. We find that AF2 does not accurately recapitulate the helix geometries in two of these proteins (**Fig. 1E, F**). Moreover, we find that the AF2 predictions for each of the proteins are systematically closer to the idealized starting geometry of 2LV8 than to the experimentally determined structure (**Fig. 1G**). Interestingly, we observe that this behavior is common to other structure prediction methods such as ESMFold^4^, OmegaFold^26^, and RGN2^27^ (**Supplementary Fig. 2**).

To test whether this prediction bias towards idealized protein geometries is more general than observed in the two examples above, we next generated 10,000 backbones with diverse geometries (**Fig. 1A**) for the 2x2 Rossmann fold using LUCS^22^. We then used ProteinMPNN, previously shown to lead to high design success rates^9^, to design sequences for each backbone. As for the examples above (**Fig. 1E, F**), we find that for designs on these 10,000 LUCS-generated backbones both AF2 and AF3 systematically predict more idealized geometries for the LHL elements than present in the LUCS design models (**Fig. 1G, Supplementary Fig. 3**A; data points above the diagonal indicate predictions closer to the idealized 2LV8 backbone than the LUCS-generated design backbone). These results indicate a potential general bias of deep-learning-based structure prediction methods such as AF2 and AF3 (if run without multiple sequence alignments that are not available for de novo proteins) towards idealized geometries.

We previously showed on a small test set of 45 designed proteins that LUCS designs with diversified LHL geometries had experimental success rates of 38% to be folded and stable when individually tested^22^. However, we next wanted to evaluate whether and how many of the much larger set of 10,000 designed sequences indeed folded into stable proteins. To do so, and since individual purification and biophysical characterization on such a large number of designs is prohibitive, we used a previously established yeast display assay that had been shown to allow massively parallel evaluation of the stability of thousands of de novo designed proteins^25^. In brief, the assay selects for stable proteins by displaying designed proteins on the surface of yeast, treating the population with increasing concentrations of protease, and monitoring loss of a fluorescent tag by protein cleavage as a proxy for instability, read out by deep sequencing of FACS-selected populations of stable (uncleaved) designs at increasing protease concentrations (**Fig. 2A**). Rather than ranking proteins by stability, here we used the assay conservatively to merely distinguish well-folded and stable proteins from unstable proteins, relative to two sets of negative control sequences; in the first set, we sampled from design sequences with scrambled amino acid sequences (full scramble). In the second set, we also scrambled amino acid sequences but maintained the hydrophobic-polar pattern (patterned scramble controls, see Methods). As expected, we find that there is a clear separation between full or patterned scramble negative controls and a large fraction of the designed proteins (**Fig. 2B,C. Supplementary Fig. 4**). We set a conservative threshold to define a protein as stable if it resists higher protease concentrations than the 95th percentile of all patterned scrambled controls to two different proteases, trypsin and chymotrypsin. By this metric, 5,996 of the 10,000 de novo designed proteins were designated as stable. This success rate of 60% is higher than that previously reported using Rosetta design methods on LUCS-generated backbones (38%), presumably because of the increased design fidelity of ProteinMPNN relative to Rosetta design^9^. Despite this increase in success rate, we believe our stable design set still represents a conservative estimate as individual tests of selected designs indicate that designs below the 95th percentile cutoff also express well (**Supplementary Fig. 5**).

**Figure 2.**
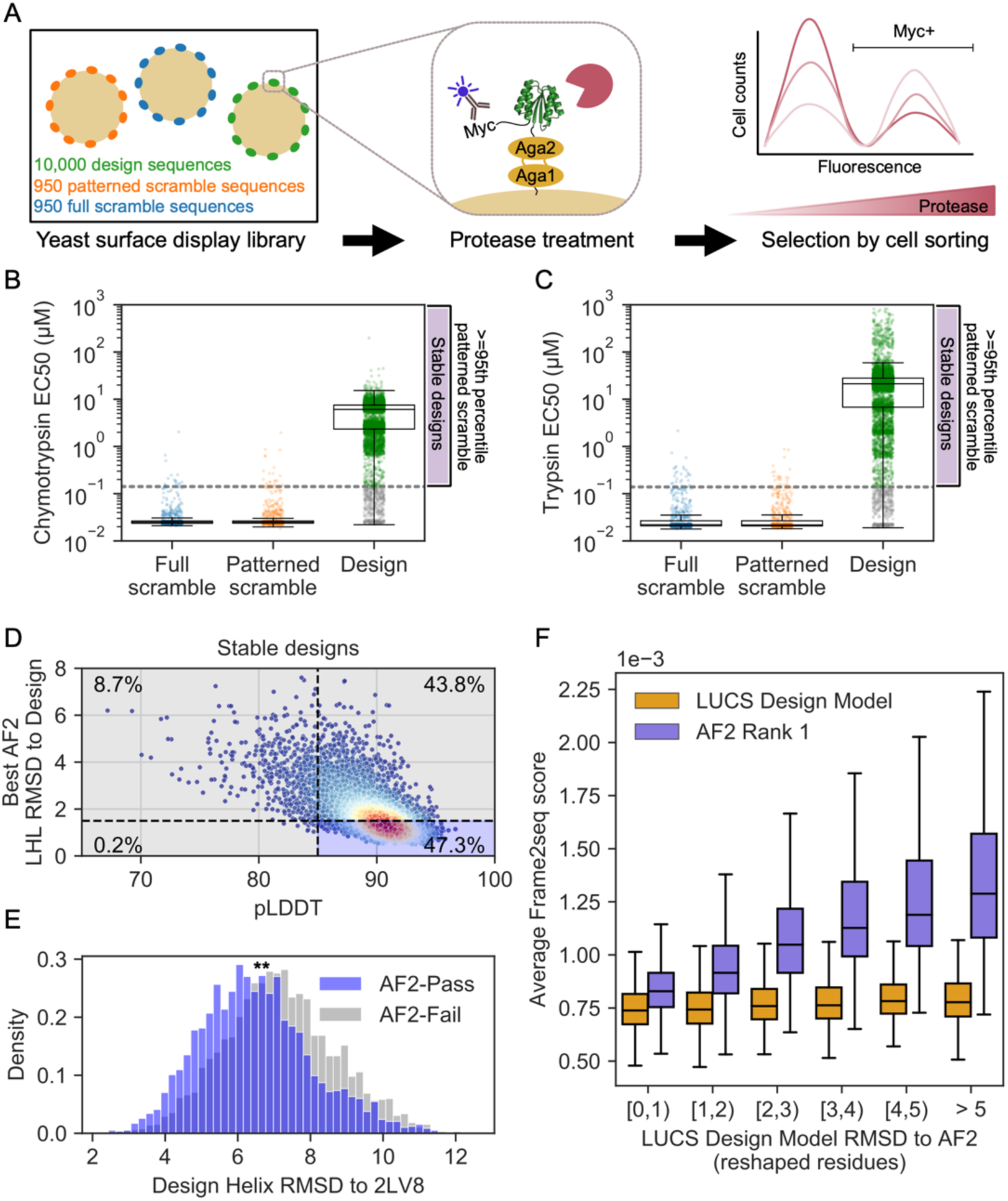
AF2 fails to capture geometric diversity in a large dataset of ∼6000 experimentally tested, stable proteins. (**A**) Schematic of yeast display assay^25^ to estimate stability of 10,000 LUCS designs along with scrambled sequence negative controls. (**B**) Boxplot with overlaid data points for resistance to chymotrypsin and (**C**) trypsin (estimated EC50s, Methods) as a metric for stability. Design sequences with EC50 for both proteases greater than the 95th percentile of patterned scramble sequences (“stable designs”) shown in green, with remaining design sequences with EC50 for one or both proteases lower than the cutoff shown in gray. Boxes show the lower and upper quartiles, whiskers extend to points within 1.5 interquartile ranges of the lower and upper quartiles, and lines indicate the median values. (**D**) AF2 metrics for stable designs, colored by density. Percentage of designs relative to typical filter cutoffs (RMSD < 1.5 Å to the design model and pLDDT > 85) labeled in each quadrant. (**E**) Distribution of helix RMSDs of stable designs that pass or fail both AF2 filters shows stable designs that fail AF2 filters have significantly higher helix RMSDs to the idealized Rossmann fold (p<0.0001 by two-sided t-test). (**F**) Frame2seq^10^ scores (negative pseudo log-likelihoods) estimating the sequence-structure compatibility of a provided backbone and sequence, binned by RMSD over the reshaped region between the LUCS design model and rank 1 (highest-pLDDT) AF2 prediction. Boxes show the lower and upper quartiles, whiskers extend to points within 1.5 interquartile ranges of the lower and upper quartiles, and lines indicate the median values (outliers not shown). Frame2seq estimates a higher sequence-structure compatibility (i.e. lower Frame2seq score) for the design model than the rank 1 AF2 prediction.

When we predicted the structures of this stable design set with AF2, we observed that many predicted structures deviated considerably from the LUCS design models, as shown for the entire set above (**Fig. 1G**). Moreover, the AF2 predictions were systematically biased towards the idealized 2LV8 backbone (**Supplementary Fig. 3**B; 32.6% of AF2-predicted helices are predicted more than 1 Å closer to 2LV8 than to the LUCS design model). We next compared the set of de novo designs identified as stable by our high-throughput assay to those that would typically have been tested experimentally if using deep learning-based design workflows. Commonly, AF2-based filters only include designs in experimental tests whose AF2 prediction has an RMSD of < 1.5 Å to the design model and a pLDDT of > 85. We applied this RMSD filter over the reshaped LHL residues, as this was the only region whose backbone varied between designs (Methods). Only 2,835 designs (47.3% of the stable designs) pass these filters (**Fig. 2D**). Moreover, the set of stable designs not passing the AF2 filters has LHL geometries that are considerably more diverse than the set of stable designs that passed the AF2 filters (**Fig. 2E)**. In summary, applying standard AF2 filters would have eliminated over 50% of the stable designs and reduced the geometric diversity of the design models (**Fig. 2E, Supplementary Fig. 3**).

The geometric diversity analyzed above is based on the original backbones (generated by the physics-based LUCS method) that were used as input for ProteinMPNN-based design. Since the AF2 structure predictions for these designs did not capture the geometric diversity of the designed backbones in many cases, we next wanted to assess the confidence in each backbone model. While we cannot determine protein structures at the scale of thousands of designs, we sought to evaluate the likelihood of each experimentally tested sequence to be better-accommodated by either the AF2 prediction or the LUCS-generated design model. To do so, we used an orthogonal method not used in any of our designs, the structure-conditioned masked language model Frame2seq^10^ (Methods). We find that Frame2seq estimates a higher sequence-structure compatibility of the design sequence given the design backbone than it does for the design sequence given the AF2 prediction (**Fig. 2F**). Moreover, the further the AF2 prediction deviates from the original LUCS design backbone, the worse the compatibility between the AF2-predicted backbone and the designed sequence becomes. In contrast, all original LUCS backbone models have high compatibility with their designed sequences. These analyses suggest that a considerable number of stable proteins (43.8%) are predicted with high AF2 confidence (pLDDT) but with structures divergent from the design models, and these AF2 predictions have low sequence-structure compatibility scored by Frame2seq.

To improve structure prediction methods for geometrically diverse proteins, we next fine-tuned AF2 on LUCS design models and sequences, following a procedure outlined in ref.^28^ (**Fig. 3A**, **Supplementary Fig. 6**, Methods). We trained three fine-tuned AF2 models. The first model was trained on stable designs, split into training and test sets using a structure-based split in which no structure in the test set was closer than 2 Å in either helix in the reshaped LHL elements to any structure in the training set (Stable Structure-Split). The second model used all stable designs for training (Stable), and the third model was trained on all designs, irrespective of whether they were classified as stable or unstable in the yeast display assay (Stable + Unstable). As expected, fine-tuning on the structure-based split of Rossmann fold LUCS designs dramatically increases the fraction of designs in the Rossmann fold test set predicted to within 1.5 Å RMSD of the design model (from 56.6% with AF2 to 97% with fine-tuned AF2, **Supplementary Fig. 6**D). These results likely indicate that the structure-based-split test set is an easy test set given the training data.

**Figure 3.**
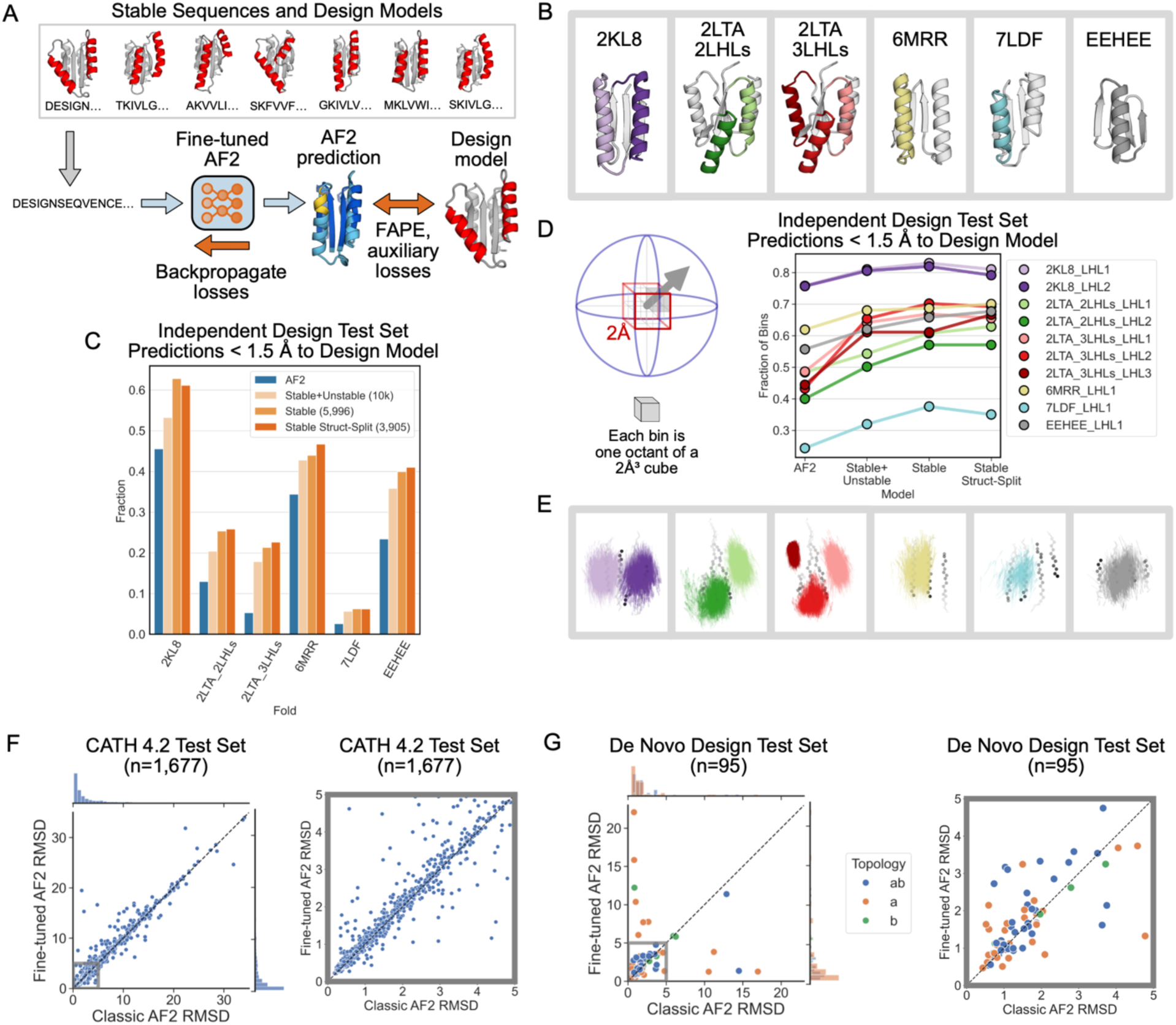
A fine-tuned AF2 model improves prediction and design of geometrically diverse proteins. (**A**) Schematic of fine-tuning AF2 using sequences and design models of LUCS designs (Methods). Three models were trained (Methods): “Stable + Unstable” (all 10,000 experimentally tested LUCS designs used in training and validation). “Stable” (5,996 stable designs used). “Stable Struct-Split” (3,905 stable designs used, structurally distinct from a test set). (**B**) Independent test set of geometrically diverse de novo proteins. Idealized starting structures are shown with colored LHL elements diversified using LUCS. (**C**) Barplot showing the fraction of well-predicted designs (< 1.5 Å RMSD to the design model) for each fold topology in the independent test set in (B) by the classic AF2 model and the three FT-AF2 models. (**D**) Geometric diversity of designs predicted well (< 1.5 Å to the design model) for classic AF2 and the three FT-AF2 models, measured by the fraction of spatial bins (cartoons on the left) occupied by designed LHL units. Cartesian and direction space are binned such that each 2 Å³ voxel is split into 8 octants^22^. (**E**) Vector representations of LUCS-reshaped helices (as in Fig. 1C) in well-predicted designs by Stable Struct-Split FT-AF2 and not the classic AF2 model. (**F**) Structure prediction performance, as evaluated by predicted versus experimental backbone RMSD, for classic AF2 and Stable Struct-Split FT-AF2 on a test set of 1,677 CATH 4.2 domains. Multiple sequence alignment (MSA) and template inputs were provided for these predictions. (**G**) Structure prediction performance for classic AF2 and Stable Struct-Split fine-tuned AF2 for a test set of 95 de novo proteins. Predictions were run in single-sequence mode (without MSA and template inputs). Right plots in (F) and (G) show closeup for low RMSD region.

To evaluate the generality of the fine-tuned model, we tested it on proteins with folds not included in the training set (non-Rossmann folds). We compiled a dataset of 5 de novo designed alpha-beta proteins with experimentally determined structures in the PDB and used LUCS to diversify the geometries of one, two, or three LHL units (generating 10,000 designs for each case; **Fig. 3B**). We then predicted the structures of the designs with both the original (classic) and the fine-tuned (FT) AF2 models. In each case, we find that the FT-AF2 models predict a larger fraction of designed sequences with an RMSD of < 1.5 Å to the design model than the classic AF2 model (**Fig. 3C**). Moreover, we find that the models trained only on the stable Rossmann fold designs (excluding unstable proteins) have increased structure prediction performance for all folds in the test set. This finding suggests that classifying sequences as stable, and training on this smaller set of designs, yields better fine-tuned model performance. For each fold, the designs predicted well (< 1.5 Å RMSD) by the model fine-tuned on the Rossman folds span a larger geometric diversity than those predicted by the classic AF2 model (**Fig. 3D,E**; Methods).

Finally, we tested whether the ability of the fine-tuned model to predict geometric diversity in de novo designed proteins degraded its structure prediction performance on natural proteins. To do so, we compiled a dataset of 1,677 CATH 4.2 domains (**Supplementary Data 2**) and used both classic AF2 and the FT-AF2 models to predict the structures using multiple sequence alignments and templates (Methods). We find that, while a few individual structure prediction results differ, the classic and the FT-AF2 models have overall similar performance (**Fig. 3F**). We also compiled a set of de novo designed proteins (**Supplementary Data 3**) and predicted their structures (without multiple sequence alignments and templates). We again find overall similar performance for the classic and FT-AF2 models (**Fig. 3G, Supplementary Fig. 7**). We did not expect better performance of the FT-AF2 model for these structures with mostly idealized geometries, as many were selected for experimental characterization because they were well-predicted by the classic AF2 model. Nevertheless, taken together the FT-AF2 models improve the prediction of geometric diversity of de novo designed alpha-beta proteins considerably including for folds unseen in training, without “catastrophic forgetting” for other proteins.

## DISCUSSION

Deviations from idealized protein geometries may be required for advanced functions such as controllable dynamics^29^ and specific small molecule binding^15^. A key finding of our study is that deep learning-based structure prediction methods such as AF2 and AF3 do not always capture the tunable geometric diversity of stable, de novo designed proteins. To propose a potential solution, we demonstrate that fine-tuning AF2 with experimentally validated and diverse stable designs improves prediction of the geometric diversity of de novo designed alpha-beta proteins deviating from an idealized starting point. In addition to the fine-tuned model, we describe a dataset of nearly 6,000 de novo designed proteins with diverse geometries that could be useful for training advanced models with better ability to capture these differences.

Our findings suggest that structure prediction models, at least in some cases, fail to capture the underlying physics of atom-level interactions, such as the subtle differences in packing interactions that specify the different geometries observed in experimentally characterized designs^22^. For natural proteins, it has been shown previously that AF2 is not sensitive to point mutations^30^. AF2’s predictions for natural proteins rely on coevolution information from MSAs, as covarying sequence positions may suggest contacts in the folded structure. A point mutation in a natural sequence will yield a nearly identical MSA to that for the wildtype sequence, and thus AF2’s predicted structure for the point mutant will be nearly identical to that of the wildtype. For de novo proteins, which have no homologous sequences and are thus predicted without coevolution information from MSAs or structural information from homologous templates, our findings indicate that AF2’s learned sequence-to-structure mapping does not accurately capture the subtle structural differences between sequences that maintain a fold topology but adopt different geometries within the fold.

We note several limitations of our study. First, we cannot experimentally determine atomic-resolution protein structures for the thousands of diverse designed proteins described here. Nevertheless, our conclusions are supported by a smaller test set of de novo designed proteins for which we do have experimental structures^22^, newly collected large-scale stability data on 10,000 designs with diverse geometries using an established assay^25^, and evaluation using orthogonal methods not trained on any of our structures and sequences^10^.

Second, while we provide experimental measurements for 10,000 de novo designed proteins, many more datasets such as the one we present here would be desirable to better assess the generality of our findings and to train better models. For instance, such datasets could include a larger diversity of folds than studied here, as well as oligomeric structures with non-idealized geometries, such that multimer models could be trained. We believe that with the rapid development of quantitative high-throughput approaches for protein characterization^31,32^ and the increasing success rates of design even for challenging engineering problems, these data will become available.

Our study provides an example approach to using informative experimental data that highlight current shortcomings to improve current deep learning models. Our results point to several areas, as outlined by the limitations above, where further informative datasets at scale are urgently needed. Ultimately, we hope that approaches such as ours will lead to improved models that allow reasoning over the detailed atom-level interactions that underlie the diverse geometries of proteins observed in nature. Such models, which should reflect more of the physical principles of atomic packing and hydrogen bonding interactions, might lead to improved generalization to more challenging design problems. It will be particularly exciting to develop and apply such models to design the non-ideal and strained geometries that make up high-energy intermediates of molecular machines, which should ultimately enable design of non-equilibrium processes driven by energy expenditure.

## SUPPLEMENTARY FIGURES

**Supplementary Figure 1.**
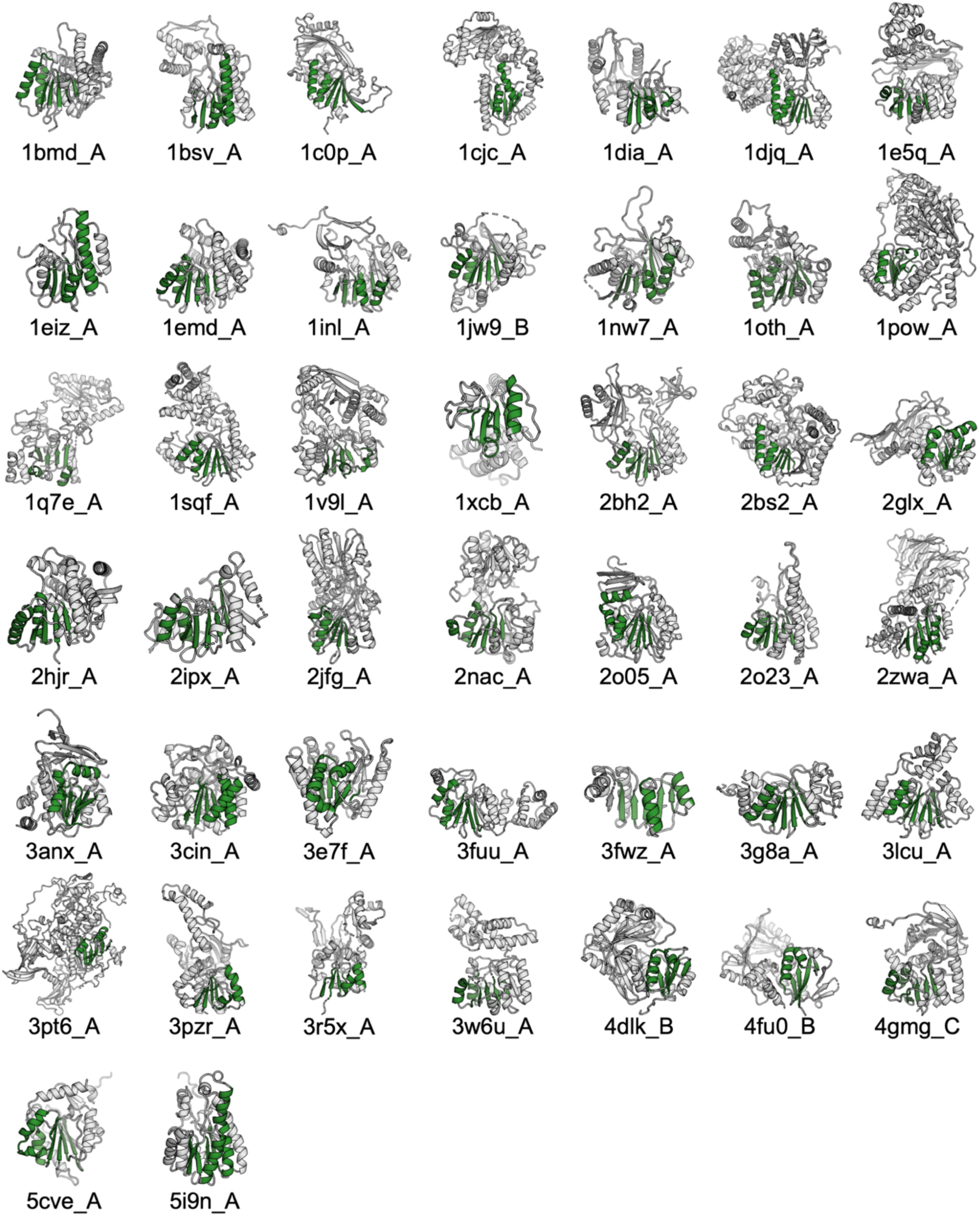
44 natural members of the Rossmann fold family. 44 natural members of the Rossmann fold family were taken from ref.^33^. Rossmann fold beta strands and topology-matched helices to the two reshaped LHLs in 2LV8 are colored green.

**Supplementary Figure 2.**
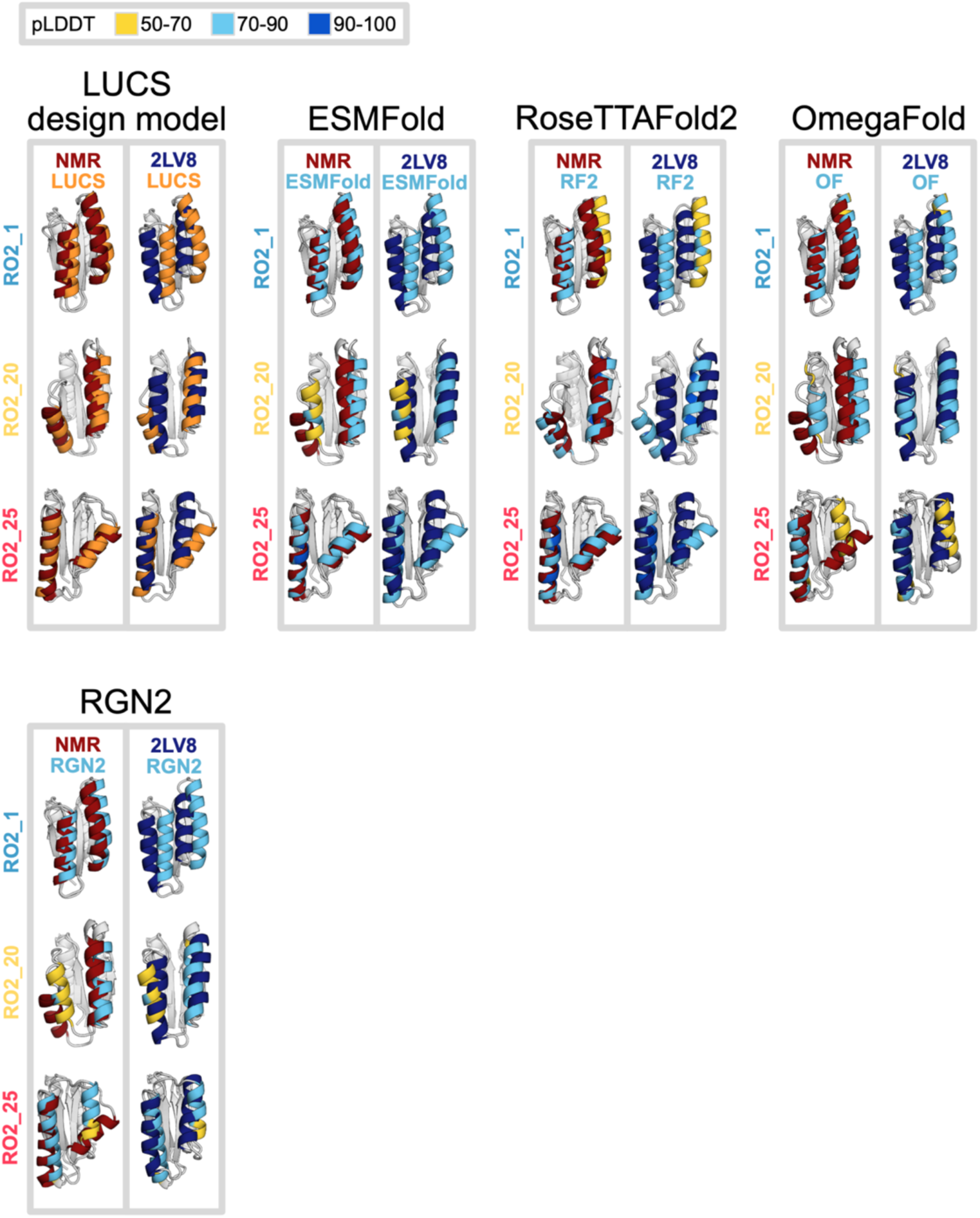
**Structure prediction for three LUCS designs with experimentally determined NMR structures using ESMFold, RoseTTAFold2 (RF2), OmegaFold (OF), and RGN2**. The LUCS design models for these three designs are also shown. Predicted structures are compared to their lowest-energy NMR structures and 2LV8 (an idealized, de novo designed 2x2 Rossmann fold protein). NMR structures are shown in dark red, 2LV8 is shown in dark blue, the LUCS design model is shown in orange, and the predicted structures are colored by pLDDT for the given prediction method. AlphaFold3^2^ (AF3) was excluded from this analysis as it currently does not support excluding template inputs of homologous protein structures. As the NMR structures of the three LUCS designs are deposited in the PDB, AF3 is able to take the ground truth structures of these sequences as inputs during inference.

**Supplementary Figure 3.**
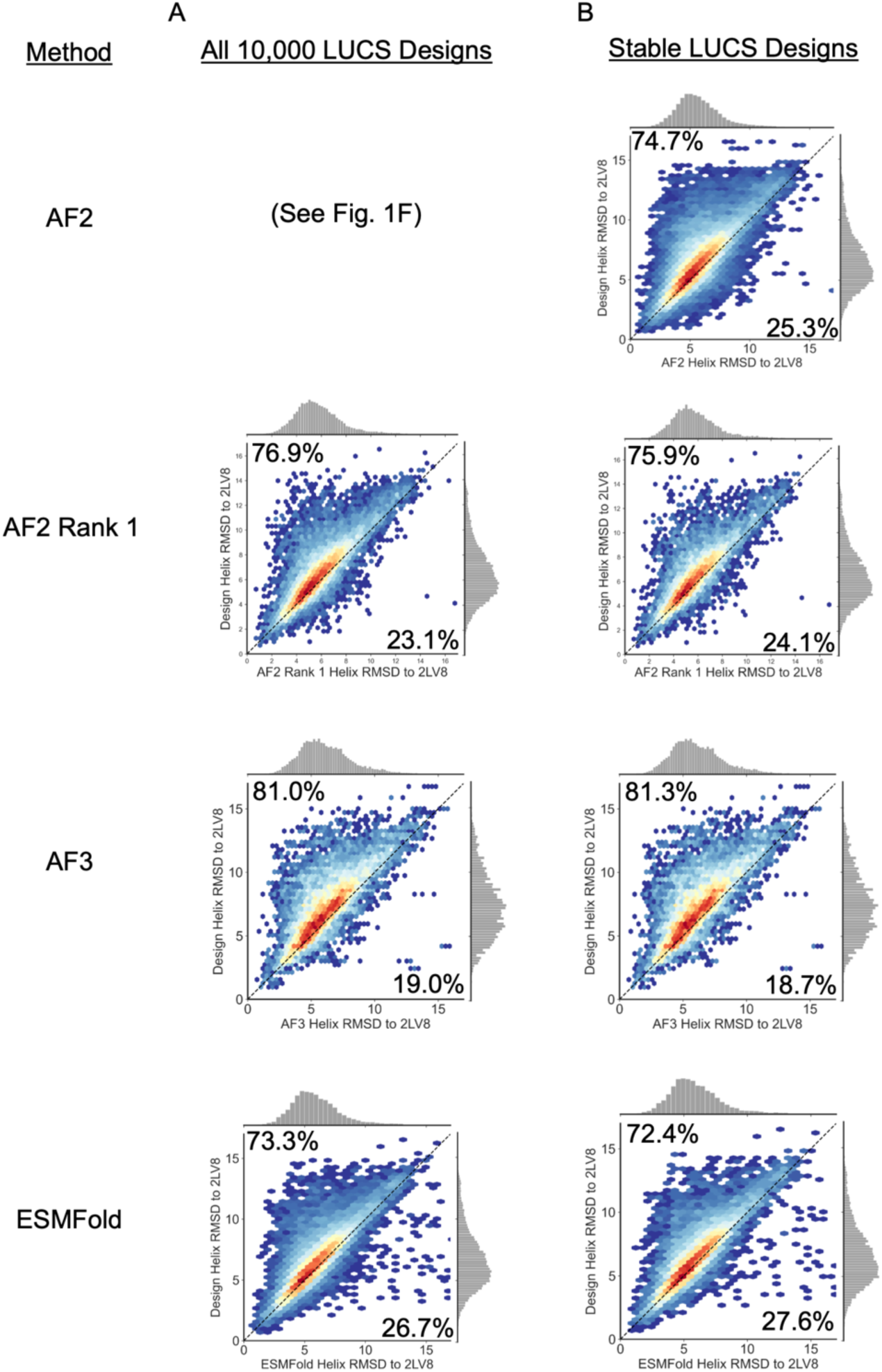
Structure prediction of Rossmann fold LUCS designs using deep learning-based protein structure prediction methods. Shown are comparisons of RMSDs for the two reshaped LUCS helices to 2LV8 for the LUCS design model (y axis) and the predicted structure from each model (x axis) for: (**A**) The 10,000 experimentally tested Rossmann fold LUCS designs and (**B**) the 5,996 stable Rossmann fold LUCS designs. All the tested structure prediction methods show a substantial bias towards the more idealized 2LV8 structure (density above diagonal, with percentage of designs indicated).

**Supplementary Figure 4.**
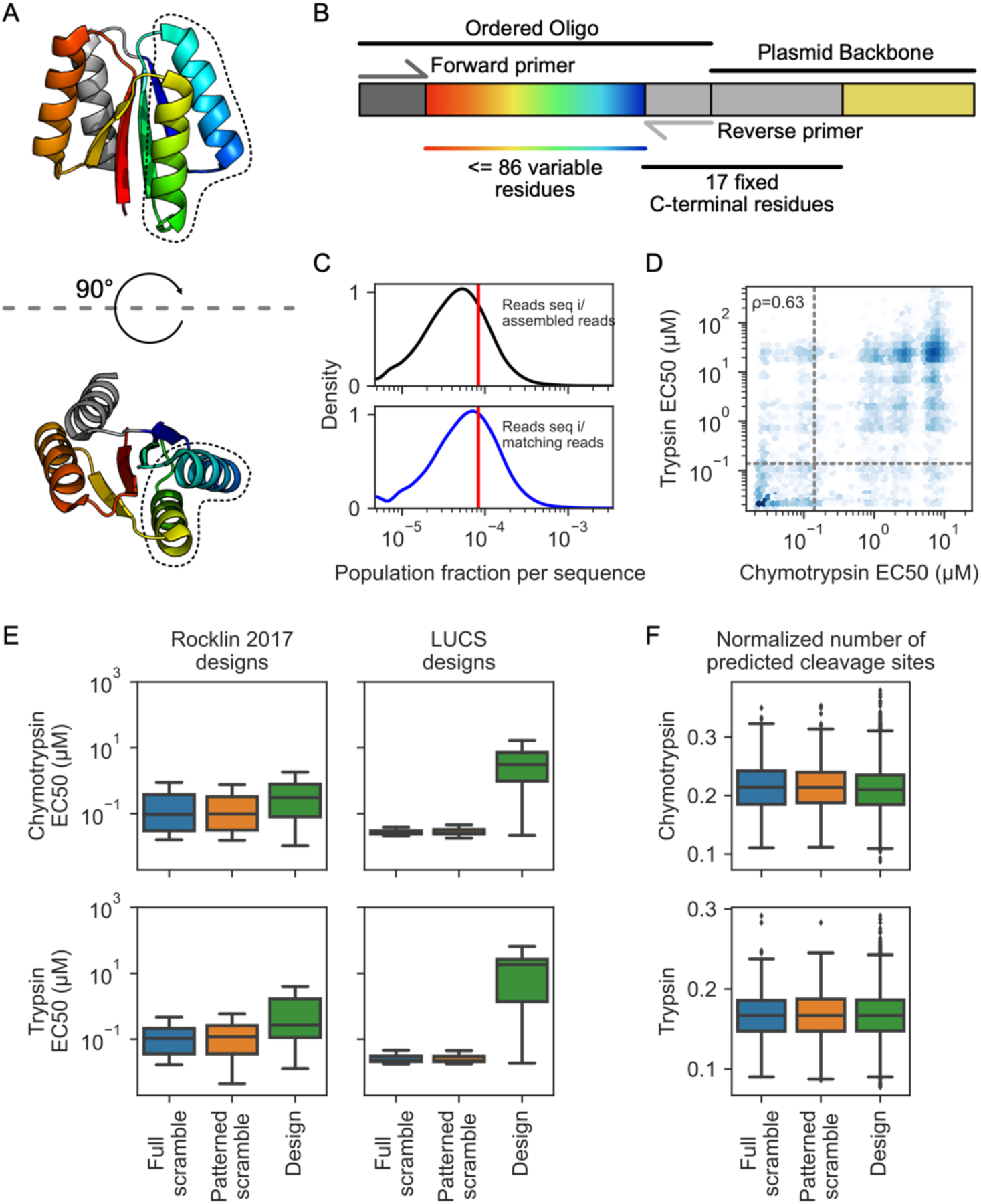
Detailed analysis of yeast display. (**A**) De novo Rossmann fold (PDB: 2LV8) showing reshaped helices on the front side of the protein (black dotted outline) and residue positions redesigned during sequence design (rainbow, red to blue representing N-term to C-term). Residue identities kept fixed during sequence redesign due to oligo pool length restrictions (300 bp) shown in gray. (**B**) Oligos were ordered with 21 base pairs on the 5’ (forward primer) and 3’ (reverse primer) ends with homology to the destination vector for library amplification and cloning by yeast assembly. This left up to 258 variable nucleotides (which encode up to 86 variable residues, shown in rainbow). The reverse primer region encodes the first 7 amino acids of the 17 C-terminal residues that were kept fixed in all designs (gray), with the plasmid backbone encoding the remaining 10 fixed C-terminal residues, which enabled testing designs with up to 103 residues in total. (**C**) Number of reads corresponding to ordered designs in the naive library expressed as a population fraction of all assembled reads (black, top panel) or assembled reads matching ordered sequences (blue, bottom panel). The expected population fraction for the ordered library of 12,000 sequences shown as a red line. (**D**) Estimated chymotrypsin and trypsin EC50s for all sequences are modestly correlated by Spearman rho (ρ=0.63). Dotted lines represent the 95th percentile of patterned scramble sequences used as a cutoff for stable designs, such that stable designs are found in the upper right quadrant. (**E**) Estimated chymotrypsin and trypsin EC50s for all sequences tested here (LUCS designs) or previously designed de novo miniproteins^25^. Negative controls (fully scrambled sequences and patterned scramble sequences – scrambled sequences that maintain the original sequence’s hydrophobic-polar pattern) show a clear separation from the designed sequences, with a much greater observed separation between controls and designs for LUCS designed proteins than de novo miniproteins. (**F**) Predicted number of chymotrypsin (top panel) and trypsin (bottom panel) cleavage sites across scramble controls and ordered designs by Rapid Peptides Generator^34^ are similar between scramble controls and designs.

**Supplementary Figure 5.**
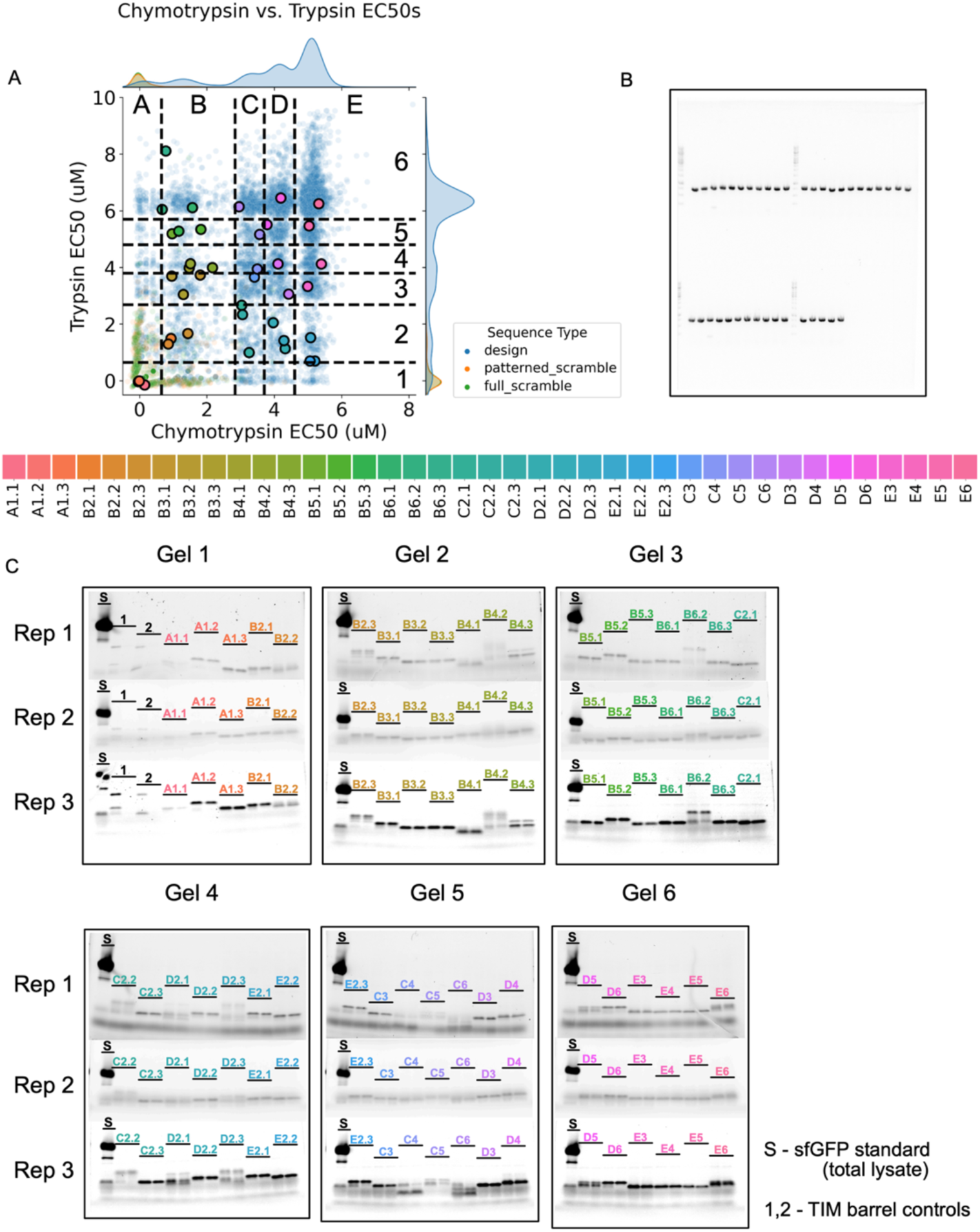
Expression tests for 39 Rossmann fold LUCS designs. (**A**) 39 designs were sampled from chymotrypsin and trypsin EC50 bins. For each bin, the design with the median helix RMSD to 2LV8 was selected. For bins in which three designs were sampled, the designs at the 25th and 75th percentiles for helix RMSD to 2LV8 were also selected. (**B**) In-gel fluorescence shows successful PCR amplification for all 39 genes. (**C**) Protein expression and solubility were tested using a cell-free protein synthesis and solubility assay (Methods). Gels show fluorescence emission at 500 nm. Each pair of lanes for each sample contains the total lysate (left) and soluble fraction (right). The total lysate of a positive control gene, encoding Superfolder GFP (sfGFP)^35^, was used as a standard for each gel. Controls 1 and 2 were de novo designed TIM barrels, which express but are not present in the soluble fraction.

**Supplementary Figure 6.**
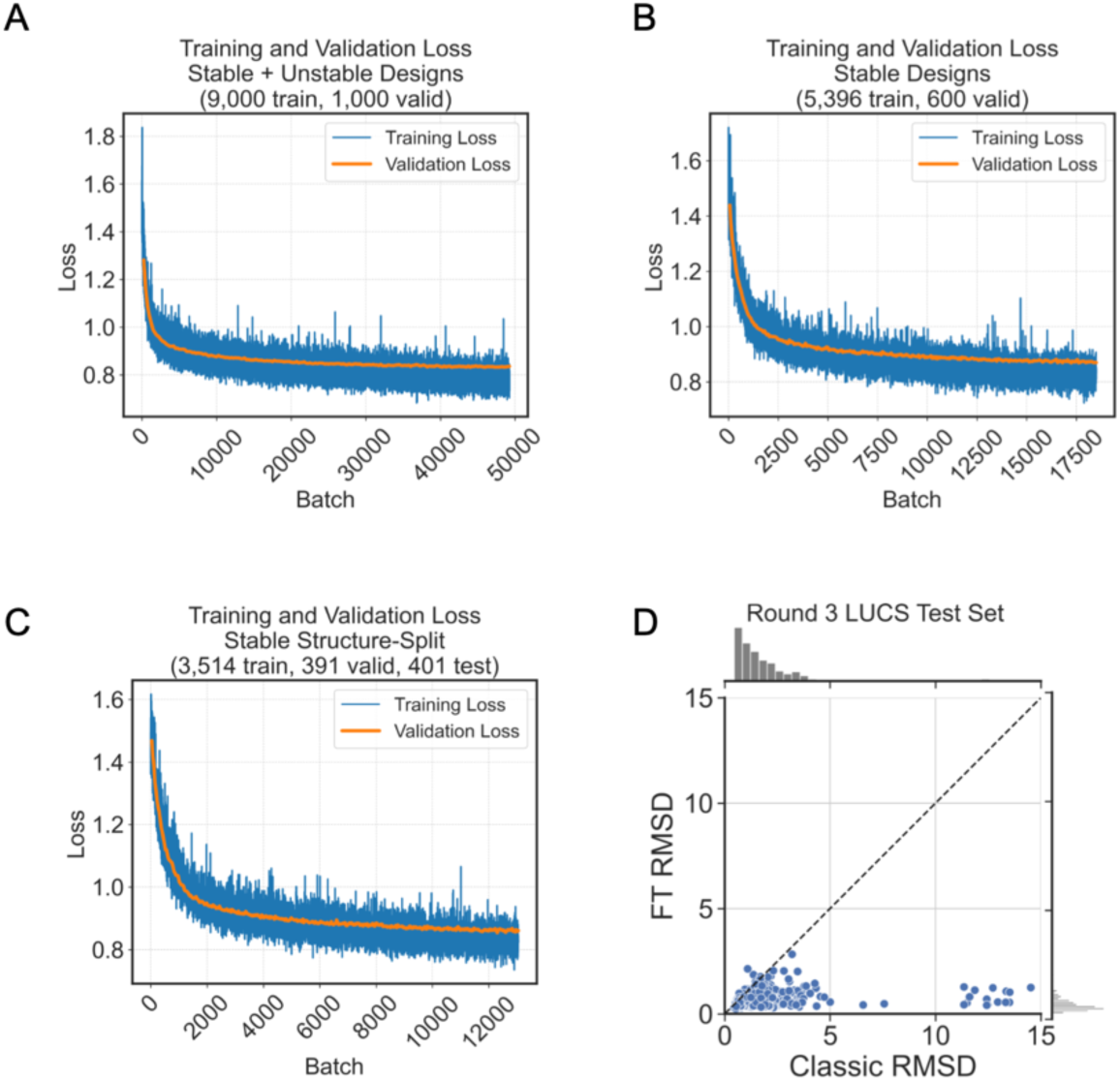
Details for fine-tuning AF2 on LUCS design test set. Training (blue) and validation (orange) loss curves for three fine-tuned AF2 models: (**A**) Stable + Unstable, (**B**) Stable, and (**C**) Stable Structure-Split. (**D**) Performance of Stable Structure-Split FT-AF2 on its test set, generated using a structure-based split in which no test set example had a helix RMSD of 2 Å or less to any topology-matched reshaped helix in the training or validation set (Methods). Stable Structure-Split greatly outperforms classic AF2 in predicting these test set examples, which likely indicates that this test set is an easy test set given the training data.

**Supplementary Figure 7.**
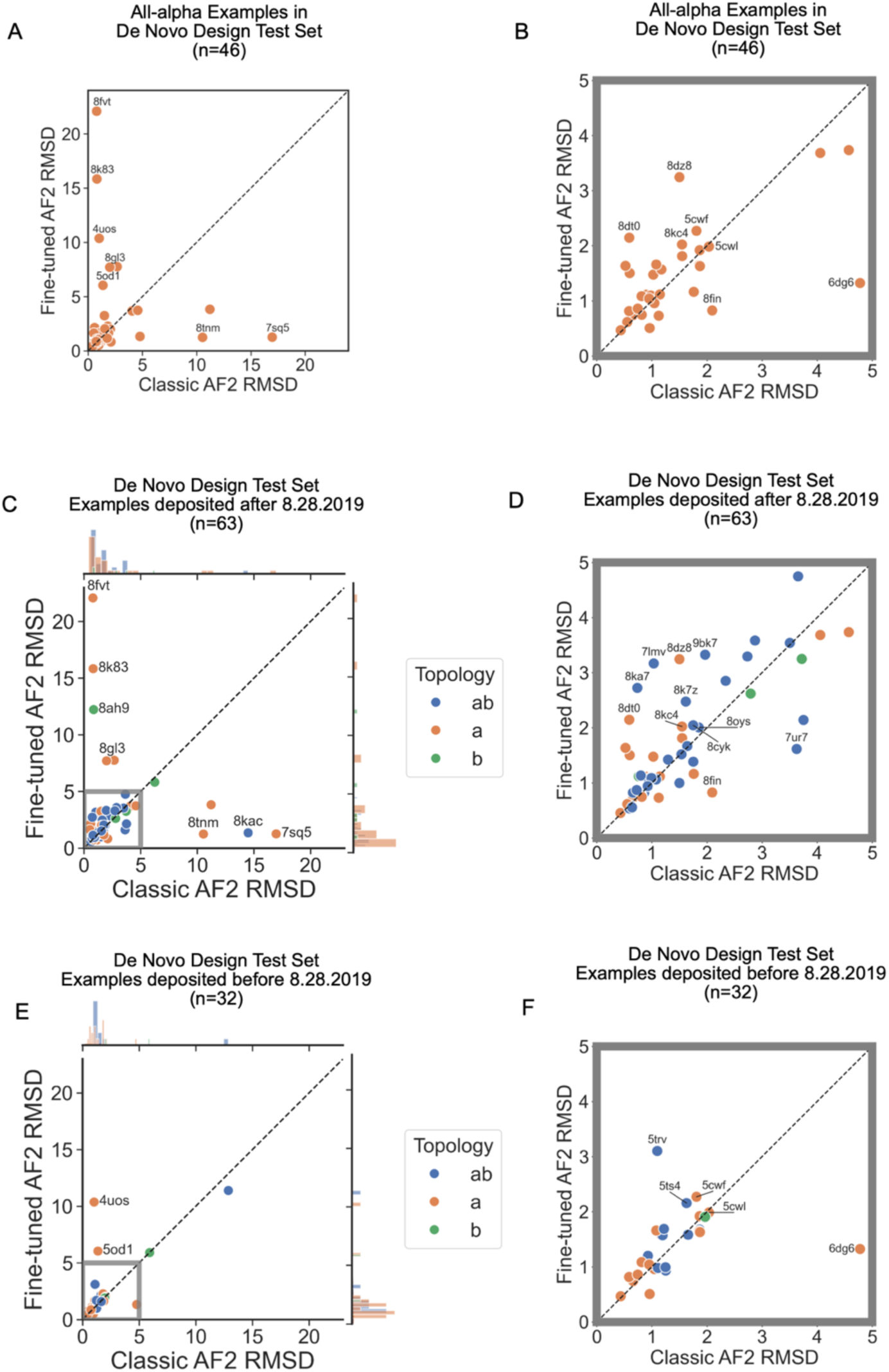
Details of predictions for de novo test set. **(A)** Predictions for the 46 all-alpha proteins in the de novo protein test set by Classic and Stable Structure-Split FT-AF2. Axes limits trimmed to 5 Å in (**B**). (**C**) De novo test set filtered by PDB deposition date after the AF2 training set cutoff (August 28^th^, 2019). Axes limits trimmed to 5 Å in (**D**). (**E**) De novo test set filtered by PDB deposition date before the AF2 training set cutoff (August 28^th^, 2019). Axes limits trimmed to 5 Å in (**F**). Examples predicted to < 2 Å RMSD to the ground truth by one model and not the other are labeled with their PDB IDs.

## METHODS

### Rossman fold dataset

The 44 natural Rossmann fold proteins (Fig. 1A, Supplementary Fig. 1) were taken from ref.^33^ This set was selected to be representative of the diversity of natural Rossmann fold protein cofactor binding functions. Rossmann fold beta strand and helix residues were manually annotated and matched topologically to the 2x2 Rossmann fold LUCS designs.

### LUCS structure generation for experimentally tested designs

LUCS was run on the PDB ID: 2LV8 backbone as previously described^22^, reshaping the loop-helix-loop elements between residues 31-51 (LHL1) and 58-76 (LHL2). There were 37,886 reshaped LHL1 and 34,986 reshaped LHL2 backbones found not to clash with the rest of the 2LV8 backbone (using poly-Valine sequences in these residue stretches to assess clashes). 50 million pairs of reshaped LHLs (3.846% of all 1.325 billion possible combinations) were sampled and screened to find compatible pairs of reshaped LHL backbones that do not clash (again, with poly-Valine sequences). 5.027 million (10.055%) of these 50 million sampled pairs of LHLs did not clash with each other. We removed backbones with more than 103 residues (to fit gene synthesis requirements, see below) and randomly selected 22,000 backbones for further evaluation.

### ProteinMPNN sequence design for experimentally tested designs

22,000 unique Rossmann fold LUCS backbones selected from the set above were designed with ProteinMPNN using sampling temp = 0.1, generating 5 sequences for each design, yielding 110,000 sequences (ProteinMPNN model: ProteinMPNN/vanilla_model_weights/v_48_020.pt). The 17 C-terminal residues were fixed to maintain their sequence in 2LV8, VTSPDEAKRWIKEFSEE (see Methods: Experimentally Tested Design Selection).

### Generating design models for experimentally tested designs

To generate design models of ProteinMPNN sequences designed onto LUCS backbones, the ProteinMPNN sequences were threaded onto the LUCS backbones in PyRosetta^36^. These structures were then relaxed using PyRosetta FastRelax, with the Rosetta score function ref2015. PyRosetta was initialized with the following flags: init(’-use_input_sc -input_ab_scheme AHo_Scheme -ignore_unrecognized_res - ignore_zero_occupancy false -load_PDB_components false -relax:default_repeats 2 - no_fconfig -ex1 -ex2’). The threaded and relaxed Rosetta models are referred to as LUCS “design models.”

### RFdiffusion structure generation

To generate structural diversity in the same protein region as was reshaped in Rossmann fold LUCS backbones, RFdiffusion backbone generation was run as a motif scaffolding problem, in which the 2LV8 backbone, excluding the two diversified LHL units, was treated as a motif to keep fixed (in LUCS, the input protein backbone, aside from the reshaped LHLs, is kept fixed). To control for LHL length effects on helix diversity, each RFdiffusion backbone generation task was run with specified lengths of diffused residues at the LHL1 and LHL2 positions that exactly matched the lengths of the pairs of LHLs in the stable LUCS designs (using the following arguments: inference.input_pdb=2lv8.pdb contigmap.contigs=[A1-29/’$LHL1_length’/A51- 56/’$LHL2_length’/A76-98]). In this fashion, RFdiffusion was used to generate reshaped LHLs that were length-matched to each of the LHLs in the 5,996 stable LUCS designs. Any RFdiffusion-generated backbone that contained fewer than 7 helical residues at either reshaped LHL unit was removed from the analysis in Fig. 1A, as was its corresponding length-matched LUCS backbone.

### Analysis of structural variation using helix RMSD

Pairwise helix RMSDs were calculated between each pair of Rossmann fold backbones within each group (natural, LUCS, and RFdiffusion). Each pair of Rossmann folds was aligned on their beta sheet residues, and RMSDs were calculated over the two reshaped helices of each pair (or, in the case of the natural Rossmann folds, the topology-matched helices to the reshaped helices in the LUCS and RFdiffusion groups). In cases where pairs of backbones contained helices of different lengths, helix RMSDs were calculated between the longest common helices (taking the central helix residues of the longer helix and including the N-terminal residue in cases where an odd number of residues were trimmed). For both the natural Rossmann fold proteins and RFdiffusion-generated backbones, helix residues were determined by DSSP. For the design models of experimentally tested LUCS designs (see Methods: Experimentally Tested Design Selection), helix residues were determined from the linker lengths used in generating the diversified LHL geometries.

### Calculating LHL RMSD between AF2 predictions and design models

AF2 predictions were aligned to design models by the non-reshaped LHL residues, and RMSD was calculated between the LHL residues. The lowest LHL RMSD of the 5 predicted AF2 models was taken for analysis in Fig. 2D.

### AF2 structure prediction

For all de novo designed proteins (the Rossmann fold LUCS designs and the test set of other alpha-beta LUCS designs), AF2 predictions were run using localcolabfold^37^ in single-sequence mode (no MSA or template inputs) and with 24 max recycling steps.

### Structure prediction with other methods (ESMFold, AF3, RoseTTAFold2, OmegaFold, RGN2)

For RoseTTAFold2^3^, MSA and template inputs were excluded (as is common practice for predicting the structures of de novo designed sequences, as these sequences have no homologous sequences). Additionally, for models which employ recycling steps (RoseTTAFold2^3^ and OmegaFold^27^), inference was run with 24 max recycling steps, except for AF3^2^, which was run with 3 recycling steps.

### Experimentally Tested Design Selection

10,000 unique Rossmann fold LUCS backbones, designed with ProteinMPNN, were selected for experimental testing as detailed in the sections below.

*Length restrictions for orderable designs*: At the time of ordering, oligo pools were restricted to sequences 300 bp or less. Experimentally tested oligos also included 21 identical nucleotides at the 5’ and 3’ ends for PCR amplification, leaving up to 258 variable nucleotides (which encode up to 86 variable amino acids). The 21 3’ nucleotides (reverse primer region) encoded the first 7 amino acids of the 17 C-terminal residues that were kept fixed in all designs, and the plasmid backbone 3’ of the reverse primer encoded the remaining 10 fixed C-terminal residues, which enabled testing designs with up to 103 residues in total (with up to 86 variable residues; Supplementary Fig. 4B).

*Filtering for orderable designs:* A pool of 110,000 ProteinMPNN-designed sequences (from 22,000 unique LUCS backbones; see Methods: LUCS structure generation for experimentally tested designs) was first filtered for containing 103 residues or less (see above). Designs containing Cysteine residues were also filtered out. 68,344 sequences from 17,708 unique backbones passed these filters.

*Filtering by computational metrics:* All 68,344 sequences were filtered by Rosetta and AF2 metrics. The Rosetta metrics were calculated on the LUCS design models for each sequence as described previously^22^ using i) fragment quality for the reshaped LHL units < 1 Å, ii) Rosetta holes scores < 0 for shell 2 residues (see Methods: Shell residue definitions), iii) Rosetta helix complementarity scores for reshaped helices > 0.6, iv) fewer than 2 buried unsatisfied hydrogen bonds for reshaped LHL residues, v) no oversaturated hydrogen bonds for shell 2 residues, and vi) the ratio of hydrophobic solvent accessible surface area (SASA) over the total SASA < 0.58 for shell 2 residues. The AF2 filters were: i) the highest pLDDT of the 5 AF2 models over the reshaped residues > 85 and ii) the lowest reshaped LHL residue RMSD between the 5 AF2 models and the design model < 1.5 Å. Reshaped LHL residue RMSDs were calculated by aligning the AF2 predictions and design models by their non-reshaped residues, then calculating the RMSDs between the reshaped LHL residues. Designs were divided into four groups: each combination of passing and failing the two sets of filtering metrics (i.e., Rosetta-Pass/AF2-Pass, Rosetta-Pass/AF2-Fail, Rosetta-Fail/AF2-Pass, Rosetta-Fail/AF2-Fail).

*Selecting 10,000 unique backbones:* We wanted to experimentally test 10,000 unique backbones, sampling 2,500 backbones from each of the four Rosetta/AF2 Pass/Fail groups. However, individual LUCS backbones had multiple ProteinMPNN sequences in the set of 68,344, which were often distributed over multiple Rosetta/AF2 Pass/Fail groups. Given that a disproportionately large number of designs failed the Rosetta filters, the two Rosetta-Pass quadrants needed to be populated first to ensure that at least 2,500 unique-backbone designs fell into each of these Rosetta-Pass groups. Unique LUCS backbones with designs in both the Rosetta-Pass/AF2-Fail and Rosetta-Pass/AF2-Pass quadrants were first evenly distributed between these two quadrants, removing any other designs from the initial pool of 68,344 that were designed onto the same backbone. The remaining unique backbones in these two Rosetta-Pass quadrants were assigned to their respective quadrants, again removing any other designs from the remaining pool that were designed onto the same backbone. Next, the Rosetta-Fail/AF2-Pass quadrant was populated with its remaining unique LUCS backbones from the now thrice-filtered pool of designs. Finally, the Rosetta-Fail/AF2-Fail quadrant was populated with all the remaining unique LUCS backbones in the pool. This resulted in each quadrant containing over 2,500 designs from unique LUCS backbones. 2,500 designs from each of these four quadrants were then randomly sampled, resulting in a set of 10,000 designs with unique LUCS backbones for experimental testing.

### LUCS design library

All designed protein sequences were ordered from Twist Biosciences as an oligo pool. Protein sequences were reverse translated and codon optimized for expression in *S. cerevisiae*. The oligo library consisted of 10,000 LUCS backbone+ProteinMPNN sequence designs (see Methods: Experimentally Tested Design Selection) and 1,900 scramble negative control sequences. For the negative control sequences, 950 sequences were sampled from the pool of 10,000 designed sequences such that 238 sequences were randomly sampled from each of the Rosetta-Pass/AF2-Fail and Rosetta-Fail/AF2-Pass groups of 2,500 designs, and 237 sequences were randomly sampled from the Rosetta-Pass/AF2-Pass and Rosetta-Fail/AF2-Fail groups of 2,500 designs. Two types of control scramble sequences were generated: “full scramble” controls randomly shuffled all amino acid identities in the sequence, and “patterned scramble” controls maintained the hydrophobic-polar pattern of the amino acids in the sequence but shuffled identities within these two groups while keeping Gly and Pro residues fixed.

### Library amplification and yeast transformation

Libraries were amplified using KAPA HiFi polymerase (Roche #07958927001) in two rounds of PCR (12 cycles, each round) using the following primers:

Forward: 5’-GGAGGCTCTGGTGGAGGAGGTAGTGGAGGTGGAGGCAGTGCTTCAATG-3’

Reverse: 5’-CCTCACTGAATTCTTTAATCCAACGCTTAGCCTCGTCCGGAGAAGTCAC-3’

The gel extracted product of the first round PCR was used as input template for the second round PCR. Gel extracted products of the second round PCR were used for yeast transformation.

Linearized yeast display entry vector pETconV4, a gift from David Baker, modified for increased protease resistance, was generated by restriction digest with SalI (New England Biolabs, #R0138S) overnight at 37°C. NdeI and BamHI (New England Biolabs, #R0111S, R3136S) were added to the reaction the next day and digested overnight at 37°C, followed by a final one hour digest at 37°C with fresh SalI, NdeI, and BamHI added to ensure complete digestion of the backbone.

*S. cerevisiae* (ATCC MYA-4941DQ, strain EBY100) was thawed from a glycerol stock by streaking out on yeast extract peptone dextrose (YPD) agar. YPD liquid culture + penicillin-streptomycin (PS) (Gibco #15140122) was inoculated with an isolated colony and cultured at 30°C, 270 rpm overnight. 25 mL YPD was inoculated with the overnight culture to a final OD600=0.2 and grown to OD600=1.5. 20 mL of culture was pelleted (∼3x10^8^ cells) and resuspended in 5 mL 100 mM lithium acetate (LiAC) and 10 mM dithiothreitol (DTT), followed by shaking at 30°C, 180 rpm for 10 min. Yeast were pelleted and washed with 2.5 mL sterile ice-cold water. Yeast were pelleted and resuspended in 100 μL sterile ice cold water. Half of competent yeast (∼1.5x10^8^ cells) were transformed with 2.5 μg of the amplified library and 1 μg linearized pETcon-v4 vector and half with 1 μg linearized vector alone as a negative control. Yeast were electroporated using the BTX Gemini X2 Twin Wave electroporation system (BTX #452040) at 500 V, 15 ms single pulse in a 2 mm gap cuvette and recovered in YPD at 30°C for 1 hr without shaking. Transformation efficiency was assessed by serial dilution plating of the library and control transformations on SD-CAA plates to ensure at least 10x coverage of the library and that the negative control transformation has a transformation efficiency < 1% of the library transformation. Isolated colonies were sent for Sanger sequencing to confirm correct cloning of the library via yeast recombination.

### High-throughput yeast protein stability assay

Protein stability was assessed using a high-throughput yeast display protease susceptibility assay as described in ref.^25^. Briefly, library expression was induced by culturing overnight at 30°C, 270 rpm in SG-CAA medium. Yeast were digested with either Trypsin (Gibco #15-400-054) in phosphate-buffered saline (PBS) pH 7.4 or Chymotrypsin (MilliporeSigma #C4129) in Tris-buffered saline (TBS)+100 mM CaCl2 pH 8.0 for 5 min at room temperature before quenching with ice cold PBS+1% bovine serum albumin (BSA) or TBS+1% BSA and subsequently washed three times before labeling with 1:100 anti-Myc Alexa Fluor 647 conjugate (Cell Signaling Technologies, #2233S) for 15 min at room temperature. Yeast were washed twice and resuspended in chilled PBS+BSA and stored on ice until sorting.

Yeast were sorted on the BD FACSAria III using purity settings. APC gate for the Myc-positive population was set on an uninduced, protease untreated, anti-Myc-labeled population. 10x library coverage of the Myc-positive population was collected for all protease treatment conditions. Sorted yeast were recovered for 30 min at 30°C in YPD medium, before growth overnight in SD-CAA. The next day a minimum of 1e7 cells were pelleted and frozen at -80°C for DNA extraction and the remaining culture used for the next round of protease digestion where applicable. In total, three rounds of sorting and selection were performed with increasing protease concentrations in each round with the cells treated with the highest concentration of each protease in each round used as the input for the next round of sorting (Trypsin, round 1: 0.07 μM, 0.21 μM; round 2: 0.64 μM, 1.93 μM; round 3: 5.78 μM, 17.33 μM, 51.99 μM. Chymotrypsin, round 1: 0.08 μM, 0.25 μM; round 2: 0.74 μM, 2.22 μM; round 3: 6.67 μM, 20 μΜ.)

### DNA extraction and NGS

Plasmid DNA was extracted from yeast using the Zymoprep Yeast Plasmid Miniprep II kit (Zymo Research Corporation, #D2004). The entire elution was used as input template for PCR amplification using KAPA HiFi polymerase (Roche #07958927001) to introduce partial Illumina adapter sequences for Amplicon-EZ sequencing with Azenta.

Forward: 5’-ACACTCTTTCCCTACACGACGCTCTTCCGATCTGGAGGTGGAGGCAGTGCTTC-3’

Reverse: 5’-GACTGGAGTTCAGACGTGTGCTCTTCCGATCTGTCCTCTTCAGAAATAAGCTGTTGT TCG-3’

Amplified products were run on an agarose gel to confirm the major PCR product was the amplicon of the expected size. The remaining reaction was purified by PCR cleanup (Qiagen #28104) and sent for amplicon sequencing at AZENTA, Inc. (South Plainfield, NJ, USA). Library preparation and sequencing were performed using the NEBNext Ultra DNA Library Prep kit following manufacturer recommendations (Illumina, San Diego, CA, USA). Samples were sequenced using 2x 250 paired-end configuration. Raw sequencing data was demultiplexed using bcl2fastq version 2.17.1.14. Trimmomatic version 0.36 was used to trim adapter sequences and filter low quality basecalls, and reads less than 30 base pairs long were discarded.

### EC50 from sequencing counts

Analysis of raw sequencing data was carried out using the scripts from the following repository: https://github.com/Haddox/prot_stab_analysis_pipeline as described in brief below with minor modifications.

Briefly, reads were paired using the PEAR program and a read considered a count for a given sequence if, 1) it contained the six bases immediately preceding the design coding sequence at the 5’ end (TCAATG) and the six bases immediately following the design coding sequence, including the fixed C-terminal 17 amino acids, at the 3’ end (CTCGAG), and 2) matched the complete amino acid sequence of the ordered design.

Protease resistance was determined using the previously reported model^25^, which assumes that proteolysis follows pseudo-first order kinetics with a rate constant specific to each sequence. Designs were included in downstream stability analyses if their EC50 95% credible interval < 2.0 (equivalent to two selection rounds) (6,748 / 10,000 experimentally tested designs had ec50_95ci < 2.0) as described previously^25^.

We used EC50 directly as a metric for protein stability, rather than previously reported stability score. We reasoned that since we observe a clear difference in EC50 values for scrambled controls versus designs that EC50 likely adequately captures foldedness of our protein fold of interest, unlike what is observed for previously characterized miniproteins^25^ (de novo designs and controls from rounds 1-4) which are smaller in size and may be more susceptible to sequence-dependent effects (Supplementary Fig. 4E). Further, the predicted number of chymotrypsin (high and low specificity) and trypsin cleavage sites based on sequence using Rapid Peptides Generator^34^ shows a similar number of predicted sites across controls and designs, suggesting the large differences in EC50s measured for scramble controls vs designs are unlikely to be explained by the number of cleavage sites in different sequences.

### Small-scale protein expression tests

*Sampling 39 designs:* We selected 39 designs for small-scale experimental expression tests. Designs were sampled from 2-dimensional bins defined by Chymotrypsin EC50 and Trypsin EC50 thresholds (**Supplementary Figure 5A**). For bins with three designs sampled, design with the median helix RMSD to 2LV8 was sampled, as were the designs at the 25th and 75th percentiles for helix RMSD to 2LV8. For bins with 1 design sampled, the design with the median helix RMSD to 2LV8 was sampled.

*Gene amplification:* Linear DNA templates containing design sequences were amplified using PCR. PCR products were visualized by DNA gel electrophoresis and fluorescence imaging (**Supplementary Figure 5B**).

Each 25 μL PCR reaction contained:

- 12.5 μL 2x Q5 Hot Start High-Fidelity 2X Master Mix (New England Biolabs, Product Number: M0494S)
- 1.25 μL Forward Primer (from 10 μM stock)
- 1.25 μL Reverse Primer (from 10 μM stock)
- 2.5 μL DNA template (from 1 ng/μL stock)
- 7.5 μL water

PCR cycling protocol:

1. 30 seconds at 98°C
2. 10 seconds at 98°C
3. 30 seconds at 63°C
4. 42.5 seconds at 72°C
5. 120 seconds at 72°C
6. Hold at 12°C

A 120 mL agarose (0.8% weight/volume) gel was prepared by adding:

- 120 mL Tris-Acetate-EDTA (TAE) buffer
- 0.96 g agarose
- 12 μL GelRed Nucleic Acid Gel Stain (Biotium, Product Number: 41003)

2 μL of each of the 39 PCR products were mixed with 1 μL of Gel Loading Dye, Purple (6X) (New England Biolabs, Product Number: B7024S) and 3 μL of water. 5 μL of this mixture was loaded into each gel lane. 5 μL of GeneRuler 1 kb DNA ladder was also loaded. The DNA gel was run at 110 V and 100 mA for 45 min. The gel was imaged using Bio-Rad Gel Doc EZ imager (Bio-Rad, Product Number: 170-8270).

*Small-scale protein expression test:* Protein expression tests were performed using an adapted version of the protocol outlined in ref.^38^. The primary deviation from this protocol was the use of Fluorotect Dye (a BODIPY-labeled Lysine loaded onto tRNA; Promega FluoroTect™ GreenLys in vitro Translation Labeling System, Product Number: L5001) and fluorescence imaging (GE Typhoon FLA 9000 Imager) to detect protein expression instead of radiolabeled Leucine and autoradiography.

For each of the 39 designs, DNA encoding the design was added to a 1.5 mL tube (15 uL total reaction volume) along with:

- 30% (by volume) S30 E. coli lysate
- 2 mM amino acids
- 6 mM Mg(Glutamate)₂
- 10 mM ammonium acetate
- 130 mM K(Glutamate)
- 1.2 mM ATP
- 0.85 mM GTP
- 0.85 mM UTP
- 0.85 mM CTP
- 0.03 mg/mL Folinic acid
- 0.17 mg/mL tRNA
- 0.4 nM NAD
- 0.27 mM CoA
- 4 mM Oxalic acid
- 1 mM Putrescine
- 1.5 mM Spermidine
- 57 mM HEPES
- 30 mM PEP
- 13.33 ng/μL plasmid
- 2% (by volume) Fluorotect Dye
- 26% (by volume) water

This mixture was incubated for 20 hours at 30°C. During incubation, the Fluorotect Dye is incorporated into newly synthesized proteins (i.e., the tested LUCS designs) in place of their Lysine residues. After incubation, RNAse cocktail (Invitrogen™ RNase Cocktail™ Enzyme Mix) was added to each reaction to digest the Fluorotect Dye and prevent further protein synthesis. A sample of the total lysate was taken from each tube, then the tubes were spun down at 18,000 x g for 20 min. Finally, a sample was taken from the supernatant of each tube. The total lysate and supernatant samples from each tested design were run in a 4-20% Tris-glycine gel (Bio-Rad Mini-PROTEAN TGX Precast Gels, Product number: 4561091) with SDS running buffer. SDS-PAGE gels were run at 180 V and 100 mA for 42 min. Fluorescent signal at 510 nm was visualized using GE Typhoon FLA 9000 Imager (**Supplementary Figure 5C**).

### S30 extract preparation

S30 extracts were prepared from *E. coli* BL21 Star (DE3) (Thermo Fisher). Cells were streaked onto LB-Agar and incubated overnight at 37 °C. A single colony was inoculated into LB and grown overnight at 37 °C with 220 rpm shaking. 1 L of 2xYPTG was inoculated at OD_600_= 0.075 and grown at 37 °C with 220 rpm shaking. At OD_600_ = 0.6, the culture was induced for T7 RNA polymerase expression using 1 mM IPTG. The culture was harvested at OD_600_= 3.0 by centrifugation at 5k x g for 15 minutes at 4 °C. The pellet was washed three times with cold S30 buffer (10 mM Tris-Acetate pH 8.2, 14 mM Mg Acetate, 60 mM K Acetate) and stored at -80 °C.

Frozen pellets were thawed on ice and resuspended in 1 mL of S30 buffer per gram of wet cell mass. Cells were lysed with 9 cycles of 5s ON, 5s OFF sonication at 15% power. The cell lysate was centrifuged at 12,000 x g at 4 °C for 10 minutes. Supernatant was collected and spun again at 10,000 x g at 4°C for 10 minutes to ensure that insoluble components were removed. The lysate was aliquoted into 50 µL portions, frozen in a dry ice ethanol bath, and stored at -80 °C.

### Shell residue definitions

Shell 1 residues included all reshaped LHL residues, plus any residue that was i) within 10 Å (Cɑ - Cɑ) of any residue in the reshaped LHLs and ii) pointed toward any reshaped LHL residue, as previously described^22^. Shell 2 residues included all shell 1 residues, plus any residue that was i) within 8 Å (Cɑ - Cɑ) of any shell 1 residue and ii) pointed toward any shell 1 residue. This is the same definition used for “movable” residues in ref.^22^.

### Frame2seq scoring

To assess sequence-structure compatibility, we compute sequence likelihoods given structure and sequence using Frame2seq^10^. Frame2seq models learn to approximate P(sequence|structure) via a masked language modeling objective during training. At inference time, Frame2seq models can be further utilized as a scoring tool to compute fitness (or compatibility) of structure and sequence pairings. Given de novo/predicted structures and their corresponding (designed) sequences as input, we compute Frame2seq negative pseudo log likelihoods (PLL) as explored in ref.^39^. We separately score the compatibility of LUCS and AlphaFold2 backbones for the same designed sequences to output negative PLL as follows:

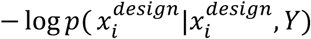

where i iterates over reshaped residue positions, x^design^ is the corresponding designed sequence, Y is structure.

### AF2 fine-tuning

Trainable AF2 code, written in JAX, was adapted from ref.^28^ for fine-tuning AF2 on LUCS designs. AF2 weights were initialized from AF2 model_2_ptm^1^. All parameters in AF2’s model_2_ptm were trainable during fine-tuning. LUCS design models were provided as the ground truth structures during training, and a random 10% of the training data was used as a validation set. For some fine-tuning experiments, a structure-based split of the experimentally tested LUCS designs were held out as a test set (see Methods: Structure-based split of Stable LUCS designs for AF2 fine-tuning). Early stopping with patience 20 was employed (i.e., model convergence was determined by when the validation loss had not decreased in 20 validation steps, at which point the model checkpoint with the lowest overall validation loss was saved as the final model). Validation loss was calculated at intervals depending on the sizes of each fine-tuned model’s training and validation sets. For the Stable Structure-Split fine-tuned AF2 model, validation loss was calculated every 600 training steps (3,514 train, 391 validation examples). For the “Stable” fine-tuned AF2 model, validation loss was calculated every 1,000 training steps (5,396 train, 600 validation examples). For the “Stable + Unstable” fine-tuned model, validation loss was calculated every 4,000 training steps (9,000 train, 1,000 validation examples). A batch size of 16 was used in all fine-tuning runs. A learning rate schedule of 0 to 0.01 was used over the first 1,000 training steps, after which the learning rate was kept at 0.01. Gradient scaling with the Adam Optimizer was used, with a decay rate for the exponentially weighted average of gradients (b1) of 0.9, a decay rate for the exponentially weighted average of squared grads (b2) of 0.95, and an epsilon term (small positive constant added to the denominator for numerical stability) of 1e-6.

### Structure-based split of Stable LUCS designs for AF2 fine-tuning

To ensure that the train and test sets were structurally dissimilar, spectral clustering was performed on stable LUCS designs and groups of clusters were sampled. First, the all-by-all Helix 1 and 2 RMSDs, as well as the overall Helix RMSDs, between all 5,996 stable, experimentally tested LUCS design models were calculated. Next, spectral clustering was performed on the all-by-all overall Helix RMSDs to cluster the designs into 25 groups. Spectral clustering includes: i) computing the similarity matrix and construct the graph Laplacian, ii) computing the eigenvectors corresponding to the smallest eigenvalues of the graph Laplacian, iii) using these eigenvectors to embed the data points into a lower-dimensional space, iv) clustering the data points in the lower-dimensional space with k-means clustering. K-means is randomly initialized, and so 1,000 random seeds were tested at this clustering step to sample more data splits. For each of the 1,000 random seeds, which yielded 1,000 different sets of 25 clusters, the clusters were randomly split into two groups 1,000 different times, and these cluster splits were scored by 1) the Helix 1 and 2 RMSDs between each pair of examples across the data split and 2) the relative sizes of the groups of clusters. In the end, 1,000,000 different groups of clusters were scored on these criteria. The highest scoring cluster split was selected, and individual examples from this split were excluded such that no pairs of examples across the data split had < 2 Å Helix 1 or 2 RMSD to each other. Designs were also removed from the train and test sets such that no pair of examples across the train-test split had identical linker backbones (sampled by LUCS when reshaping LHL elements) in their reshaped LHLs. Such examples were removed from either the train or test sets as to best approach a 9:1 train:test ratio.

### LUCS structure generation for 5 alpha-beta folds

LUCS designs were generated for five different de novo alpha-beta proteins with unique fold topologies, not including the Rossmann fold (as the 2x2 Rossmann fold was included in the fine-tuning training set). The PDB IDs of the input backbones used in this test set (and their insertion points for their LUCS-reshaped LHLs) were: 2KL8 (LHL1: 8-32, LHL2: 47-71), 2LTA (LHL1: 11-29, LHL2: 35-55, LHL3: 58-80; two sets of LUCS-reshaped 2LTA backbones were generated, one with LHLs 1 and 2 and the other with LHLs 1, 2, and 3 reshaped), 6MRR (LHL1: 38-62), 7LDF (LHL1: 9-27), EHEE_rd1_0284^25^ (LHL1: 6-25). 2,000 unique LUCS backbones were generated for each input backbone and, with two different sets of LHLs reshaped for 2LTA. This procedure yielded 12,000 unique backbones in total. ProteinMPNN (ProteinMPNN/vanilla_model_weights/v_48_020.pt) was run with a sampling temperature of 0.1, generating 5 sequences for each input backbone (60,000 LUCS designs in total). Any designs which contained at least one reshaped LHL that was moved to the opposite face of the beta sheet from its starting position were removed from the test set. This left 7,651 2KL8 designs, 8,664 6MRR designs, and 9,931 EEHEE designs. All other fold groups in the LUCS test set retained all 10,000 designs. Design models were generated for each LUCS design as in Methods: Generating design models for experimentally tested designs.

### Analysis of structural variation using geometry bins

After aligning LUCS designs on their non-reshaped residues, 6-dimensional helix vectors (3 dimensions for the helix centroid, and 3 for the helix N-to-C terminal direction) were calculated for reshaped LUCS helices, as described previously^22^. Helix centroids were determined by calculating the average position of helix backbone atoms. Helix directions were determined by calculating the average carbonyl-carbon-to-oxygen direction vectors for the helix backbone atoms. The position space was divided into 2Å³ cubes and direction space was divided into 8 octants within each cube, creating 6-dimensional bins.

### AF2 structure prediction for naturally occurring proteins using multiple sequence alignments

7,555 CATH4.2 domains were filtered for 1) unique CATH domain identifiers (6,119 domains), 2) having 200 residues or less (2,784 domains), 3) having no chain breaks (1,769 domains), and 4) having no Selenocysteine, Pyrrolysine, “ASX” (ambiguous identity: either ASN or ASP), and “UNK” (Unknown) (1,677 domains). These 1,677 CATH 4.2 domains were then used as a test set of natural proteins. The 1,677 CATH 4.2 domains (**Supplementary Data 2**) were predicted using ColabFold, loaded with Classic AF2 model_2_ptm or fine-tuned AF2 parameters, with random seed = 0, and MSA and template inputs generated by MMseqs2 and HHsearch, respectively.

### AF2 structure prediction for de novo designed protein set

A set of 95 de novo proteins was gathered from the RCSB PDB. First, 1,143 structures were gathered using the following search query:

QUERY: Structure Keywords HAS EXACT PHRASE “DE NOVO PROTEIN” AND Scientific Name of the Source Organism = “synthetic construct” AND Experimental Method = “X-RAY DIFFRACTION”

Next, proteins with fewer than 20 or greater than 200 residues were removed from the set, leaving 612 examples. This set was then filtered to exclude oligomeric proteins and any engineered natural proteins (not de novo designed), leaving a set of 95 monomeric, de novo proteins in the test set (**Supplementary Data 3**). This test set was predicted by classic and fine-tuned AF2 in single-sequence mode (without MSA and template inputs) and with 3 recycles.

## Acknowledgements

We thank Dr. Kosuke Seki for advice on cell-free expression and members of the Kortemme lab for discussion.

## Funding

Supported by NIH grant R35 GM145236 (T.K.); NSF GRFP fellowships (B.O. and D.A.); the UCSF Program for Breakthrough Biomedical Research, funded in part by the Sandler Foundation (T.K. and M.J.K.); and a UCSF MAKK Seed Award (B.O.). T.K. is a Chan Zuckerberg Biohub Investigator.

## Author contributions

B.O., S.E.C. and T.K. developed the conceptual approach. B.O. performed structural analysis, computational design, structure prediction, model fine-tuning, and cell-free expression. S.E.C. performed high-throughput yeast display experiments with assistance from E.Z.. B.O. and S.E.C. conceptualized and performed data analyses. D.A. developed and performed Frame2seq scoring. T.K. and M.J.K. provided guidance and resources. T.K., B.O. and S.E.C. wrote the manuscript with contributions from the other authors. All authors read and commented on the manuscript.

## Competing interests

Authors declare that they have no competing interests.

## Data and materials availability

Design models, predictions, computational metrics, and experimental stability data will be deposited on github and zenodo and will be available upon publication. All other relevant data are available in the main text or the supplementary materials. Rosetta source code is available from rosettacommons.org.

## Notes

### Competing Interest Statement

The authors have declared no competing interest.

